# Improving sensitivity in environmental DNA measurement by reconcentrating DNA extracts

**DOI:** 10.64898/2026.07.21.739767

**Authors:** Takashi Fukuzawa, Yanjie Zhao, Naofumi Nishizawa, Hisao Nagata, Hideyuki Doi

## Abstract

Environmental DNA (eDNA) methodology is widely applied in the biomonitoring of organisms, but it requires the target DNA to be detected in a simple, stable, and highly sensitive manner. Detection sensitivity of eDNA measurement becomes particularly critical when monitoring species present at low abundance. In this study, we aimed to improve the detection sensitivity through a method of DNA-extract reconcentration. This approach involves reconcentrating eDNA samples that were originally extracted using the widely adopted DNeasy Blood and Tissue Kit (Qiagen), utilizing the same kit’s reagents, and does not require any additional equipment or reagents. We evaluated the ability of this DNA reconcentration method using field samples including river, lake and costal marine habitats. Evaluation of this DNA reconcentration method showed that when ten conventionally extracted samples were pooled, the DNA concentration increased by approximately sevenfold, as confirmed by DNA quantification and quantitative PCR analyses, demonstrating enhanced detection sensitivity.

## 1 INTRODUCTION

Environmental DNA (eDNA) refers to a method that targets DNA fragments of biological origin present in environmental samples, such as water or soil (Goldberg et al. 2016; Ariza et al. 2023), and eDNA methods have rapidly gained popularity in a decade as a non-invasive and highly efficient techniques for biological monitoring (Taberlet et al., 2018; Deiner et al., 2021; Yang et al. 2021). Compared to traditional field surveys, eDNA-based monitoring methods offer significant reductions in labor and cost (Lugg et al. 2018), and are also effective for detecting rare or endangered species (Nagarajan et al. 2024). As a result, eDNA is being increasingly applied in a wide range of the fields, especially for nature conservation, biodiversity assessment, and invasive species management (Hering et al., 2018).

In eDNA surveys, it is essential to detect target DNA in a manner that is simple, stable, and highly sensitive. Numerous technological developments of eDNA methods have been reported with the aim of improving these aspects. Enhancing simplicity is important to facilitate the widespread adoption of this technique. For example, the eDNA methods have been developed that complete measurement in the field using mobile PCR devices (Thomas et al. 2020; Doi et al., 2015), rapid and user-friendly on-site detection methods using CRISPR-Cas13 technology (Yang et al., 2024), as well as the methods streamlining filtration and extraction using microfluidic chips (Fukuzawa et al., 2022).

Maintaining the reliability of measurement results requires improving the stability of the eDNA measurement. Accordingly, there have been advancements in preservation techniques, such as the systematic analysis of the factors (e.g., water temperature and water quality) affecting the rate of eDNA degradation (Lamb et al. 2022), the development of improved storage solutions and optimized temperature conditions to prevent DNA degradation (Lopez et. al., 2024; Sales et. al., 2019), as well as enhancements to extraction methods that stabilize extraction efficiency (Fukuzawa et al., 2023).

High detection sensitivity in eDNA measurement is particularly important for detecting the presence or absence of species and is indispensable for monitoring rare or endangered species. To achieve the higher detection sensitivity, digital PCR has been employed as a highly sensitive detection method (Doi et al., 2015), and efforts have been made to increase the volume of filtered water to maximize the recovery of target DNA (Sakata et al., 2020; Kawakami et.al., 2023). However, in natural water bodies, filter clogging caused by suspended solids and organic matter often limits the water volume that can be filtered (Liu et al., 2024). In addition, field-based filtration is constrained by time and physical workload, making efficient large-scale processing difficult (Hinlo et al., 2017). It is also widely recognized to the practical limitation to the concentration efficiency that can be achieved through a single round of filtration and extraction, as demonstrated by these and other studies (Turner et al. 2014.).

To address these challenges in improving the detection sensitivity, the present study proposes and evaluates a two-step method in which eDNA samples, once concentrated and extracted, are further re-concentrated the DNA extracts. This approach would enable a simple re-concentration process that increased the overall concentration efficiency, and as a result, the detection sensitivity could be improved.

## 2 MATERIALS AND METHODS

### 2.1 Water sampling

Water sampling was conducted from the river, lake or sea surface at the shore part of the Nakatsu River (35.531942, 139.285241), Kokai River (36.077822, 140.004244), Lake Kasumigaura (36.035112°N, 140.257929°E), Tama River (35.719792, 139.328906), Nakaminato coastal area (36.337283, 140.593853) and Sagami River (35.575099°N, 139.308802°E), We used a 0.5 L plastic cup for the water sampling.

### 2.2 DNA extraction

Filtration and extraction were conducted using two different methods. One followed the procedure described in the “Environmental DNA Sampling and Experiment Manual ver. 2.2” published by the eDNA Society (Minamoto et al., 2021), hereafter referred to as the manual method. In this method, water samples were filtered using a 0.47-micron Sterivex filter unit, and materials containing eDNA collected on the filter were extracted using the DNeasy Blood & Tissue Kit (Qiagen, Hilden, Germany).

The other method, referred to as the Biryu Chip method (hereafter, BC method, Fukuzawa et al., 2022), utilized a microfluidic device called the Biryu Chip, in which both filtration and extraction were performed within the chip itself. Because the BC method uses a small volume (20 µL) for extraction, it allows for a reduced filtration volume while maintaining a comparable concentration efficiency to that of the manual method.

In the manual method, filtration was carried out at the sampling site. The Sterivex filters were then transported to the laboratory under refrigerated conditions, frozen at –20°C upon arrival, and subsequently used for DNA extraction. The extracted solution was stored frozen until analysis. In contrast, the BC method enabled immediate extraction after water collection at the field site. The extracted samples were transported under refrigeration and stored frozen at –20°C until measurement.

### 2.3 Metabarcoding analysis

Metabarcoding analysis was conducted based on the “Environmental DNA Survey Manual ver. 2.2” published by the eDNA Society (Minamoto et al. 2021), as follows. First, a first-round PCR (1st PCR) was performed using extracted environmental DNA as the template and the fish-specific MiFish-U primer set (Miya et al., 2015). Amplification was carried out with KAPA HiFi HotStart ReadyMix (Kapa Biosystems, MA, USA). The PCR conditions were as follows: initial denaturation at 95°C for 3 minutes; 35 cycles of denaturation at 98°C for 20 seconds, annealing at 65°C for 15 seconds, and extension at 72°C for 15 seconds; followed by a final extension at 72°C for 5 minutes. The resulting amplicons were purified using AMPure XP beads (Beckman Coulter, CA, USA). In the second-round PCR (2nd PCR), indexed adapter primers were used to re-amplify the purified products and add sample-specific indices for library identification. The PCR products were checked by 1.5% agarose gel electrophoresis, and bands of the target size (∼370 bp) were excised and purified using the FastGene™ Gel/PCR Extraction Kit (Nippon Genetics, Japan). The purified library DNA was quantified using a Qubit fluorometer 2.0 (Thermo Fisher Scientific, USA), normalized, and pooled in equal concentrations. Sequencing was performed using the Illumina MiSeq platform with paired-end reads (2×150 bp).

For post-sequencing data analysis, the MiFish Pipeline provided by the Atmosphere and Ocean Research Institute, the University of Tokyo (https://mitofish.aori.u-tokyo.ac.jp/mifish/, Sato et al. 2018), was used with default pipeline settings. Specifically, adapter sequences and low-quality reads with a quality score below 20 were removed from the sequence reads, after which paired-end reads were merged and sequences within the range of 260–320 bp were extracted.

Species-level identification was assigned when the sequence similarity between the query and database sequences was ≥97%. Sequences with similarity between 90% and 97% were identified at the genus level. Sequences with similarity below 90% were classified as “unassigned.” In cases where multiple species matched with equal similarity scores, the sequence was assigned to a higher taxonomic level, typically the genus, and denoted in the format “Genus_sp.”

### 2.4 Reconcentration of DNA extracts

Reconcentration of DNA extracts was performed using the DNeasy Blood&Tissue Kit according to the following procedure. First, the sample to be reconcentrated was collected in a 5-mL tube, to which equal volumes of Buffer-AL and 99% alcohol were added and stirred. This was transferred to a spin column at 650 µL, centrifuged at 6000g for 1 minute, and the liquid transferred to the collection tube was discarded. This process was repeated until there was no more liquid in the 5-mL tube. Next, Buffer-AW1, AW2, and AE were processed according to the instructions to obtain a concentrated sample. At this time, the final Buffer-AE volume was set to 100 µL. Schematic diagram of the reconcentration procedure is shown in Figure 1.

**FIGURE 1.**
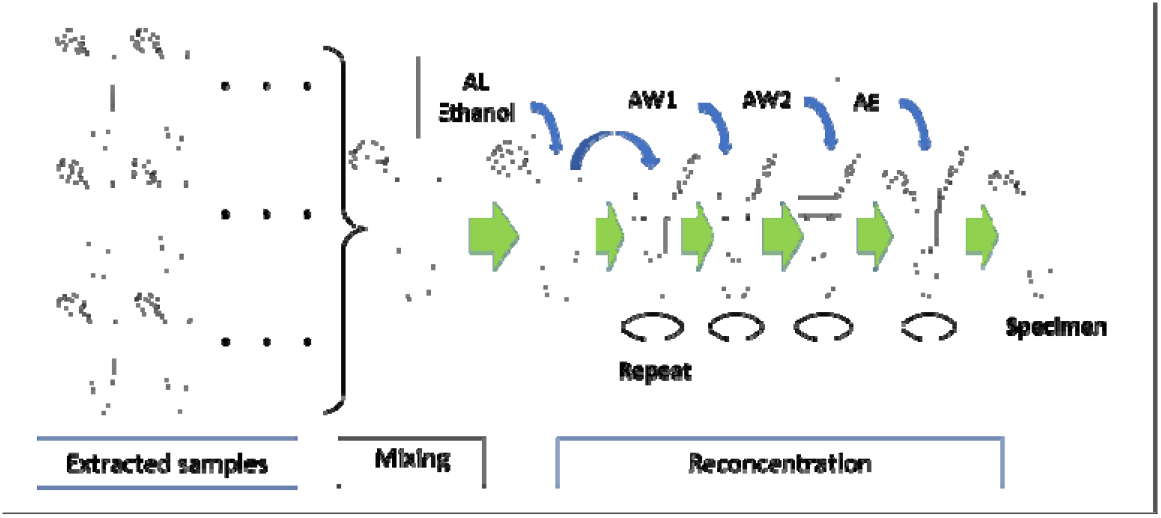
Schematic diagram of the reconcentration procedure.

### 2.5 DNA quantity measurement

Quantification of total DNA was performed using a Qubit Fluorometer and the Qubit dsDNA HS Assay Kit (Thermo Fisher Scientific, USA). The procedure followed the manufacturer’s protocol. Briefly, each sample was mixed with the Qubit working solution at a ratio of 1:199, and total DNA concentration was measured.

### 2.6 Probe-based quantitative PCR

TaqMan probe-based quantitative PCR (qPCR) was performed using a StepOnePlus Real-Time PCR System (Thermo Fisher Scientific, USA). The primer–probe sets used for the assays are listed in Table 1. Each TaqMan reaction contained of 900 nM both forward and reverse primers (Uchii et al., 2019), 125 nM of the TaqMan probe, a qPCR master mix (Easy Direct qPCR Kit [Non-treatment], Takara Bio Inc., Japan), and 1.5 µL of extracted DNA. The final reaction volume was adjusted to 15 µL with distilled water.

**TABLE 1.**
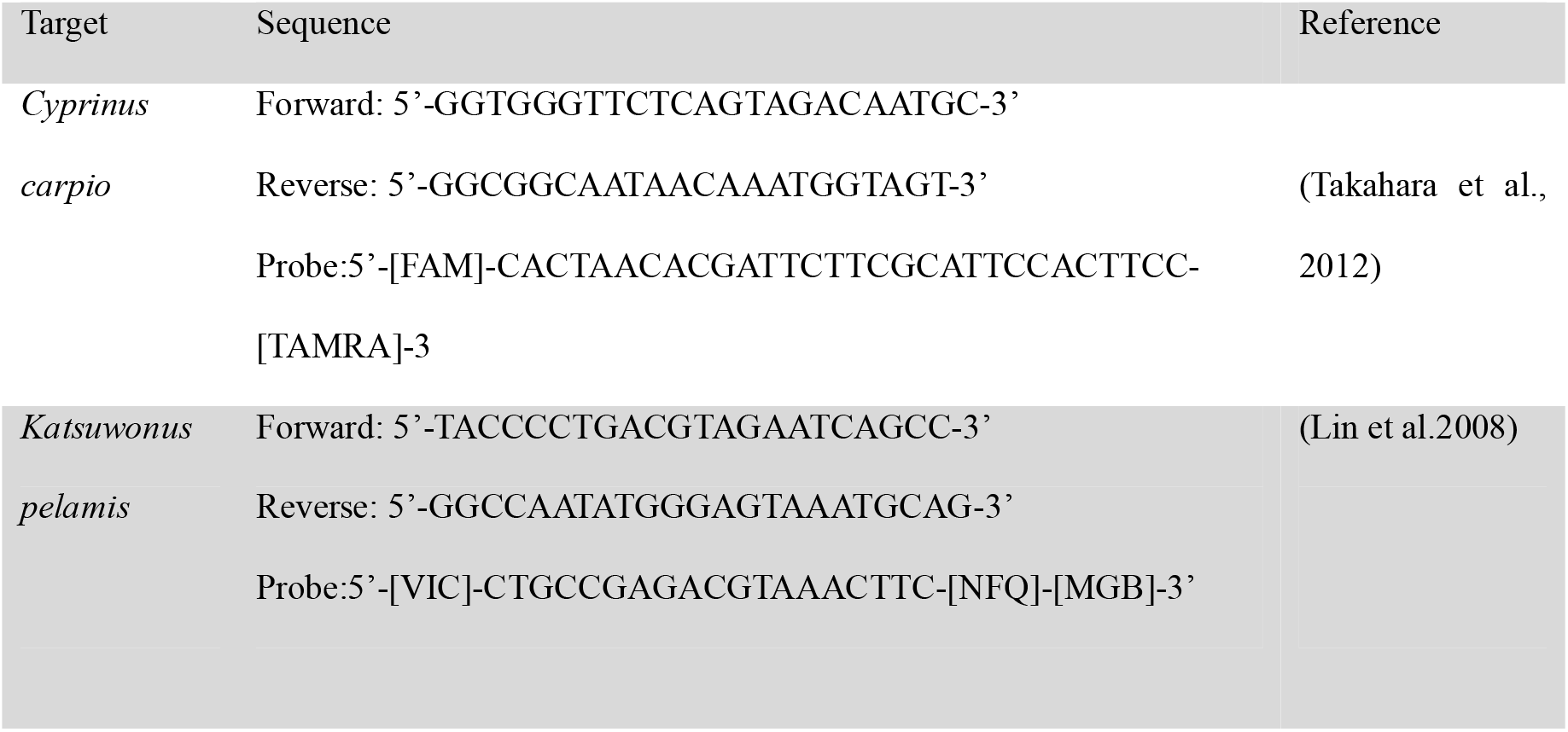
The qPCR primer-probe sets used this study.

The thermal cycling conditions were as follows: 95°C for 20 seconds, followed by 45 cycles of 95°C for 4 seconds and 60°C for 20 seconds. All reactions were performed using StepOnePlus software version 2.3.

A standard curve was generated in triplicate using a 10-fold serial dilution of standard DNA solutions at concentrations of 100,000, 10,000, 1,000, 100, and 10 copies/μL. Each sample, as well as no-template control (NTCs), was run in triplicate. The resulting standard curves had coefficients of determination (R²) ranging from 0.990 to 1.000 with PCR efficiencies between 101.0% and 102.0%.

### 2.7 Intercalator-based qPCR

Intercalator-based qPCR was performed using a StepOnePlus Real-Time PCR System. The MiFish-U primer set (Miya et al., 2015) was used for the assays. Each PCR reaction consisted of a qPCR master mix (TB Green Fast qPCR Mix; Takara Bio Inc., Tokyo, Japan), 400 nM each of forward and reverse primers (as specified in the manufacturer’s protocol), and 1.5 µL of extracted DNA. The final reaction volume was adjusted to 15 µL with distilled water.

The thermal cycling conditions were as follows: 95°C for 30 seconds, followed by 45 cycles of 95°C for 5 seconds and 65°C for 20 seconds. A melt curve analysis was performed after amplification. Fluorescence signals were recorded after holding at 95°C for 20 seconds and at 60°C for 1 minute, followed by stepwise increases to 95°C in 0.5°C increments.

For relative quantification, a three-point 10-fold serial dilution (undiluted, 1/10, and 1/100) was prepared from the sample with the highest total DNA concentration. These dilutions were used to construct a calibration curve based on the obtained Ct (cycle threshold) values. The resulting calibration curve had a coefficient of determination (R²) of 0.919 and a PCR efficiency of 96.6%.

### 2.8 Experiment 1

Water samples collected from the Nakatsu River were filtered using both the manual method and the BC method, followed by DNA extraction. For the manual method, five filtration volumes were tested: 10, 30, 100, 300, and 1000 mL. For the BC method, the filtration volume was set at 115 mL. Each filtration volume was tested in duplicate. For the extracted DNA samples, metabarcoding analysis was performed as described above, and the number of species detected was recorded.

### 2.9 Experiment 2

Water samples were collected from the Kokai River, Lake Kasumigaura, the Tama River, the Nakaminato coastal area (seawater), and the Sagami River (Table 2). At each site, six filtration replicates were performed using Sterivex filters with 200 mL of water per replicate. However, due to the high turbidity and low filterability of water at Lake Kasumigaura, the filtration volume was reduced to 100 mL. The DNA was extracted from the filtered samples using the manual method. During filtration, filter clogging typically results in increased back pressure, making it difficult to process large volumes. In this study, we defined the “filtration limit” as the volume at which the filtration rate dropped below 10 mL min^-1^ under 300 kPa of applied pressure. This threshold was used to estimate the filtration limit at each site. The water from Lake Kasumigaura exhibited particularly high turbidity, with a filtration limit of approximately 100 mL. In contrast, freshwater samples from the Kokai, Tama, and Sagami Rivers had filtration limits of approximately 500 mL, while seawater from Nakaminato site had a filtration limit of around 1000 mL.

**Table 2.** The filtration volume, filtration replicates, and Filtration limit of the samples for Experiment 2.

|  | Kokai river | Kasumigaura | Tama river | Nakaminato | Sagami river |
| --- | --- | --- | --- | --- | --- |
| Filtration Volume [mL] | 200 | 100 | 200 | 200 | 200 |
| Filtration replicates | 6 | 6 | 6 | 6 | 6 |
| Filtration limit [mL]* | 500 | 100 | 500 | 1000 | 500 |

(\* Approximate values)

At each site, six DNA extracts (200 µL each) were pooled to create a 1000 µL composite sample, with approximately 200 µL of residual extract retained separately. The composite sample (1000 µL) was subsequently subjected to a reconcentration process, and this was referred to as the “reconcentrated sample”. The remaining 200 µL of the extract, which was not subjected to reconcentration, was used as the “pre-reconcentration sample”.

For both types of the samples from each site (pre-reconcentration and reconcentrated), DNA quantification, TaqMan probe-based qPCR, and intercalator-based qPCR were conducted. In the TaqMan assay, *Katsuwonus pelamis* (skipjack tuna) was used as the target species for the seawater sample from Nakaminato coast, while *Cyprinus carpio* (common carp) was used for the freshwater samples from the other sites.

### 2.10 Statistical analysis

The analysis was performed using R version 4.3.2 (R Core Team 2023). We performed paired t-tests using the “t.test” function (paired=TRUE) to compare the differences between Before and After measurements, for each sampling site (Kokai river, Kasumigaura, Tama river, Nakaminato, and Sagami river). The dataset was first subset by site to ensure site-specific comparisons. For each site, observations with missing values (NA) were excluded from the analysis. We also tested the linear regression of ln(x) for the filtering volume and number of species detected using the “lm” function in R.

## 3 RESULTS

### 3.1 Experiment 1

Metabarcoding analyses were conducted for each sample collected from the Nakatsu River (Appendix 1). Table 3 shows the number of detected species for each sample, as well as the average number of detected species per filtration volume. The results in Table 3 indicate a general trend of increasing taxon richness with increasing filtration volume. Additionally, when comparing the manual method and the BC method, the BC method tended to detect a higher number of taxa even at the same filtration volume.

**TABLE 3.** The filtration volume and the number of species detected for each sample in Experiment 1.

| Sample |  | Manual Method |  |  |  |  |  |  |  |  |  | BC Method |  |
| --- | --- | --- | --- | --- | --- | --- | --- | --- | --- | --- | --- | --- | --- |
|  |  | 1 | 2 | 3 | 4 | 5 | 6 | 7 | 8 | 9 | 10 | 11 | 12 |
| Filtration volume [mL] |  | 10 | 10 | 30 | 30 | 100 | 100 | 300 | 300 | 1000 | 1000 | 115 | 115 |
| Concentration Ratio |  | 50 | 50 | 150 | 150 | 500 | 500 | 1500 | 1500 | 5000 | 5000 | 5044 | 5044 |
| The number of species detected |  | 9 | 9 | 10 | 7 | 9 | 17 | 16 | 19 | 20 | 20 | 20 | 22 |
|  | Average | 9 |  | 8.5 |  | 13 |  | 17.5 |  | 20 |  | 21 |  |

To evaluate these data, the following analysis was conducted. In the dataset reported by Sakata et al. (2020), the relationship between filtration volume and the mean number of detected species at three river locations (right bank, left bank, and center) followed a logarithmic pattern, as shown in Figure 2a. A logarithmic approximation provided a significant fit (R^2^ = 0.984, P < 0.0001, linear regression model).

**FIGURE 2.**
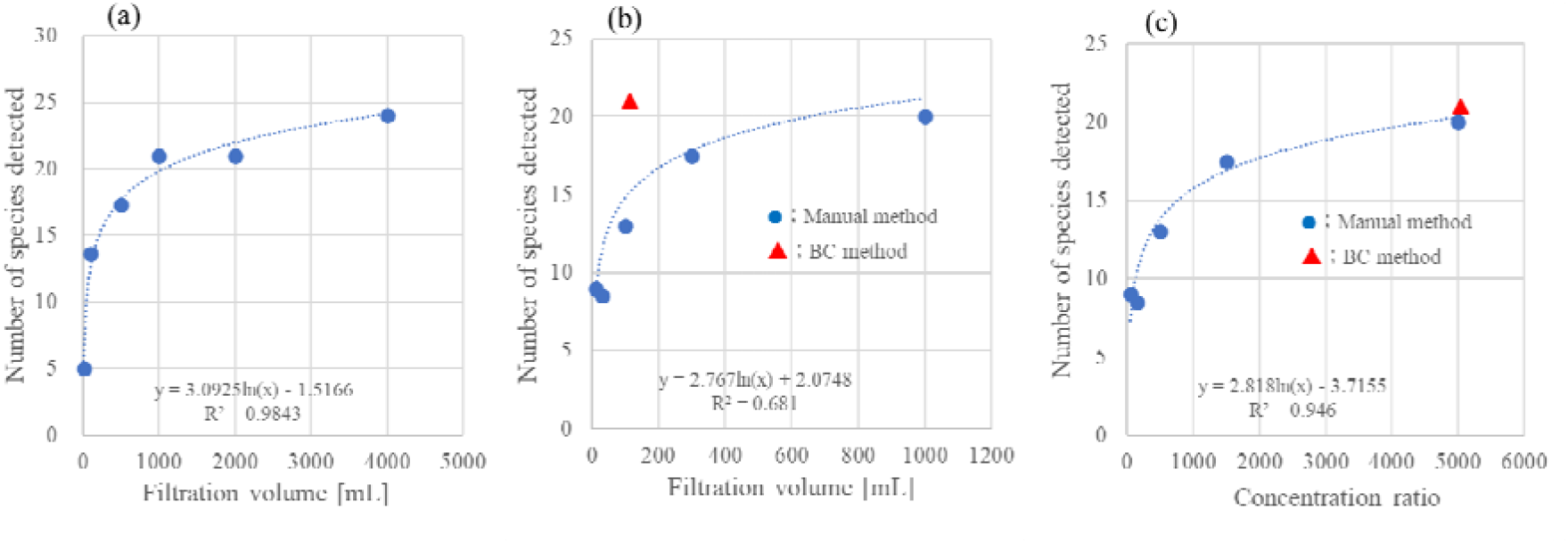
Relationship between filtration volume or concentration rate and the number of species detected. (a) Relationship with filtration volume based on the data from Sakata et al. (b) Relationship with filtration volume based on the data from Experiment 1. (c) Relationship with concentration rate based on the data from Experiment 1. The lines mean linear regression models for ln(x). When a logarithmic regression line was significantly fitted (R^2^ = 0.681, P < 0.0001, Fig. 2b, with filtration volume on the x-axis), the data points for the BC method deviated substantially above the curve. The BC method, in simple terms, maintains the concentration rate by reducing the volume of extraction solvent, allowing for reduced filtration volume while preserving detection efficiency. When concentration rate was used as the x-axis and plotted accordingly (Fig. 2c), the correlation was significantly positive (R^2^ =0.946, P < 0.0001, linear regression).

### 3.2 Experiment 2

Table 4 presents the before-and-after reconcentration results for DNA quantification, probe-based qPCR, and intercalator-based qPCR (shown as relative values only for the intercalator-based qPCR).

**TABLE 4.** DNA quantity, probe-based qPCR, and intercalator-based qPCR before and after reconcentration, obtained in Experiment 2.

| Unit | DNA quantity measurement |  |  |  | Target | Probe-based qPCR |  |  | Intercalator-based qPCR |  |  |
| --- | --- | --- | --- | --- | --- | --- | --- | --- | --- | --- | --- |
|  | Before<br>[ng/μL] | After | Total DNA<br>@after<br>[μg ] | Reconc.<br>ratio |  | Before | After | Reconc.<br>ratio | Before | After | Reconc.<br>ratio |
| Kokai river | 1.13 | 6.28 | 0.6 | 5.6 | *Cyprinus carpio* | 72.4 | 482.7 | 6.7 | - | - | - |
| Kasumi gaura | 8.90 | 75.00 | 7.5 | 8.4 | *Cyprinus carpio* | 9.8 | 59.6 | 6.1 | 28.1 | 100.0 | 3.6 |
| Tama river | 0.34 | 2.20 | 0.2 | 6.4 | *Cyprinus carpio* | 13.1 | 60.6 | 4.6 | 188.5 | 1515.2 | 8.0 |
| Naka minato | 1.09 | 8.10 | 0.8 | 7.4 | *Katsuwonus pelamis* | 0.5 | 3.4 | 6.3 | 4.8 | 51.8 | 10.7 |
| Sagami river | 0.84 | 4.50 | 0.5 | 5.3 | *Cyprinus carpio* | 5.2 | 13.5 | 2.6 | 36.1 | 229.4 | 6.3 |
| Average |  |  |  | 6.9 |  |  |  | 4.9 |  |  | 7.2 |

The t-test results (Table 5) showed most of the sites are significantly different, except in Sagami River. The reconcentration process in this study resulted in approximately a sevenfold increase in DNA concentration on average. Probe-based qPCR results also indicated that this method achieved an approximately fivefold concentration increase based on a comparison of average values. However, as shown in the individual amplification data (Appendix 2), the number of copies before reconcentration was low, leading to high variability.

**TABLE 5.** The t-test result of probe-based qPCR.

| Site | Mean before | Mean after | P Value | t Statistic | DF |
| --- | --- | --- | --- | --- | --- |
| Kokai river | 72.4 | 482.8 | 0.00214 | -21.6 | 2 |
| Kasumigaura | 9.8 | 59.6 | 0.02728 | -5.9 | 2 |
| Tama river | 13.1 | 60.6 | 0.00601 | -12.8 | 2 |
| Nakaminato | 0.5 | 3.4 | 0.03900 | -3.5 | 2 |
| Sagami river | 5.2 | 13.5 | 0.11510 | -2.7 | 2 |

Therefore, intercalator-based qPCR was performed for the samples (excluding Kokai River) in which the copy numbers before reconcentration was low in the probe-based analysis. The MiFish primers (Miya et al., 2015) were used in this assay. The results are presented as relative values, with the post-reconcentration value of the Lake Kasumigaura sample set to 100.

Since the intercalator-based qPCR can potentially amplify non-specific products, caution is necessary. However, in the present amplifications, no evidence of non-specific amplification was observed based on the first derivative of the melting curve analysis. The variation in peak melting temperatures among sampling sites was within ±1°C. This variation is considered to reflect the differences in the DNA sequences among fish species. The reconcentration method achieved a high concentration efficiency, with an average increase of 7.2-fold across all sites, including Lake Kasumigaura. It should be noted that in the case of the Lake Kasumigaura sample, DNA quantification indicated that the concentration was close to the binding capacity of the column, which may have reduced the overall concentration efficiency. The t-tests found that statistically significant differences were observed between the pre- and post-reconcentration mean values at all sampling sites (Table 6). These results confirm that the amount of DNA amplified by the MiFish primers increased significantly due to the reconcentration process.

**TABLE 6.**
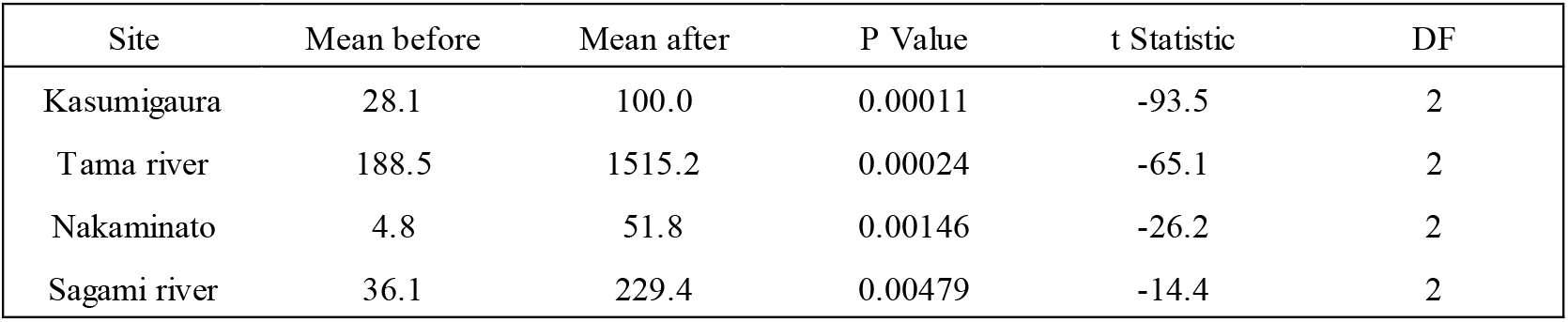
The t-test result of intercalator-based qPCR.

## 4 DISCUSSIONS

The reconcentration method developed in this study significantly enhances the eDNA detection sensitivity by increasing DNA concentration, even when the filtration volume cannot be increased. By reconcentrating ten pre-extracted samples, this approach achieved an average sevenfold increase in DNA concentration, aligning closely with the theoretical tenfold concentration rate, assuming 100% efficiency. Increasing the number of pooled samples is likely to further enhance the concentration rate. The results from Experiment 2 demonstrate that the method maintains high recovery efficiency (∼70%) when the total DNA input remains within the silica column’s adsorption capacity, as seen in the Lake Kasumigaura sample (7.5 µg) and other samples, which taking into account the filtration limits at each site, are still well below the 18 µg threshold for efficient recovery. These findings corroborate the previous studies, such as Miya et al. (2015), which emphasized the importance of optimizing DNA extraction protocols to enhance eDNA detection sensitivity in aquatic habitats.

The logarithmic relationship between filtration volume and the number of species detected in Experiment 1aligns with Sakata et al. (2020), who reported a similar pattern. However, the BC method outperformed the manual method by detecting higher species numbers at equivalent filtration volumes, suggesting that concentrating DNA through reduced elution volumes can enhance detection efficiency. This is further supported by the strong correlation between concentration rate and species detection in the BC method. These results indicate that reconcentration methods, like the one proposed here, can achieve comparable or superior detection sensitivity to increased filtration, particularly in samples with low DNA yields.

The probe-based qPCR resulted an average fivefold increase in copy numbers, highlighted the method’s effectiveness for species-specific detection, despite the variability in pre-reconcentration samples with low copy numbers, such as less than 10 copies/µL. The intercalator-based qPCR using MiFish primers demonstrated a 7.2-fold concentration increase, with consistent replicate measurements and no evidence of non-specific amplification. This suggests that the reconcentration method is particularly effective for metabarcoding applications, where amplifying target regions with universal primers is critical.

The practical applicability of the developed method lies in its simplicity and compatibility with existing laboratory equipment, such as silica columns. By concentrating 1 mL of the extract into 100 µL in this study, the method achieves significant sensitivity improvements without additional filtration, which is advantageous when filtration is limited by clogging or time constraints, particularly in turbid or marine environments (Hinlo et al., 2017; Liu et al., 2024). Further optimization, such as reducing elution volumes or increasing the volume of extract used for reconcentration, could push concentration rates closer to the theoretical maximum, provided the column’s binding capacity is not exceeded.

The results of this study demonstrate that the reconcentration method significantly enhances sensitivity in both species-specific and metabarcoding analyses. This method is straightforward to implement, utilizing existing equipment and reagents, making it a practical approach for improving detection sensitivity. Although directly not explored in this study, the washing step during reconcentration may be unnecessary, as initial sample extraction likely removes contaminants.

Additionally, replacing centrifugation with vacuum filtration for column processing could further streamline the procedure, suggesting potential for even simpler workflows.

This method offers advantages beyond improved sensitivity. By reconcentrating multiple extract prepared in parallel through filtration and extraction, it improves sensitivity and reduces processing time, particularly when filtration volumes are limited due to clogging or when seawater filtration is time-consuming. Furthermore, this approach may reduce the need for larger sample sizes to achieve higher sensitivity, allowing sequencing to be completed in a single run. This can simplify sequencing workflows and lower overall analysis costs. In applications such as multi-site or multi-timepoint measurements (e.g., daily or annual monitoring) using a pooling approach, DNA can be extracted separately and then reconcentrated collectively, offering a cost-effective method with enhanced sensitivity. While this study focused on environmental DNA analysis, the reconcentration principle may also apply to other fields, such as environmental virus detection or animal virus diagnostics, potentially enabling higher sensitivity in these contexts.

## 5 CONCLUSIONS

In this study, we here proposed a method for reconcentrating eDNA extracts intending to enhance the detection sensitivity. The reconcentration procedure can easily be carried out using existing reagents and equipment. It was found to achieve a roughly sevenfold increase in DNA concentration. These results suggest that this method enables more sensitive detection in species-specific assays and metabarcoding analyses in eDNA studies.

## Data availability

All the data are available on Supplementary Table S1 and S2.

## Acknowledgments

This study was supported by the Environment Research and Technology Development Fund (JPMEERF20204004) and JSPS KAKENHI (22K18429).

## Appendix 1

**Table S1.**
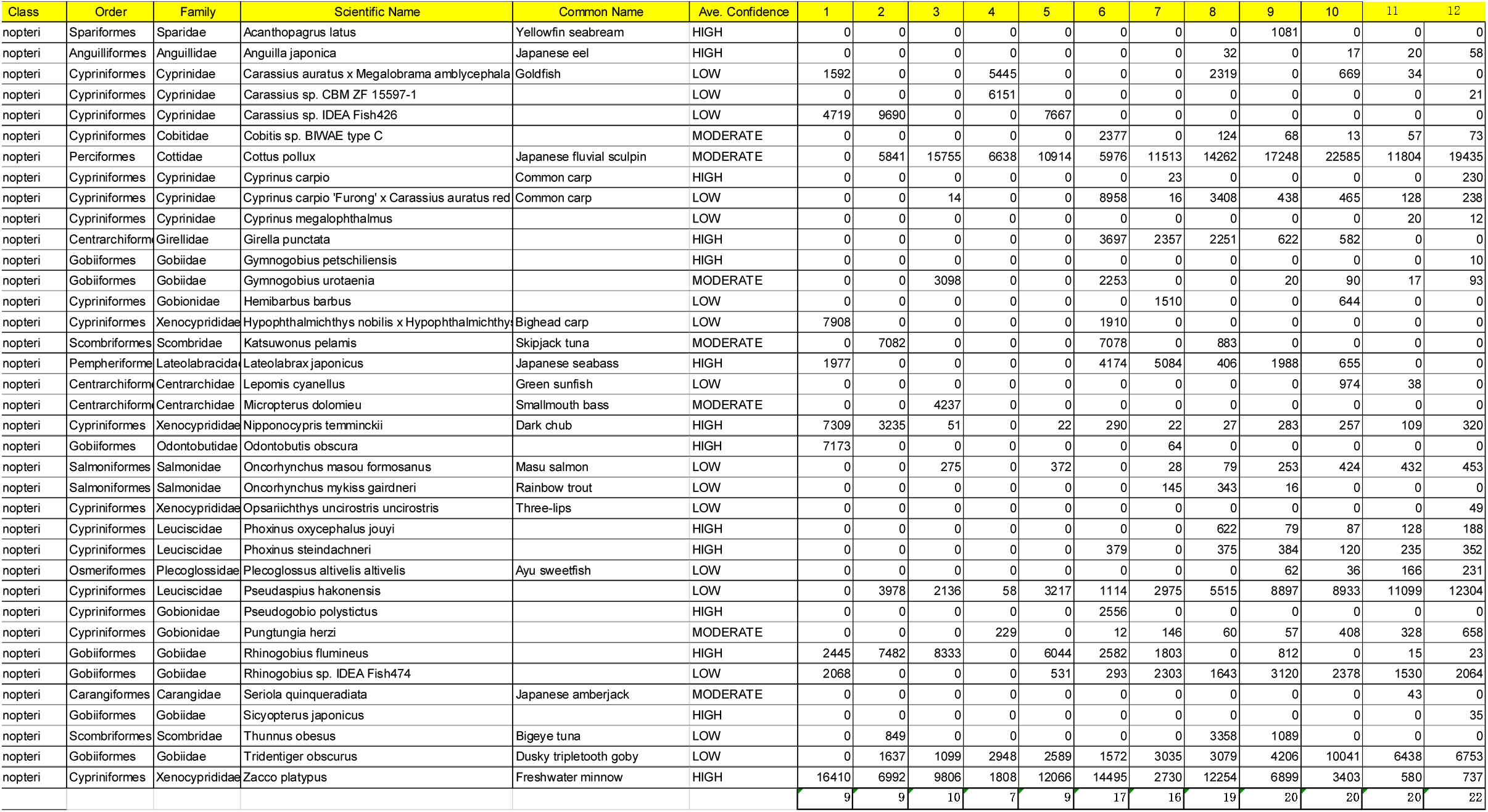
Metabarcoding results for the samples.

## Appendix 2

**Table S2.**
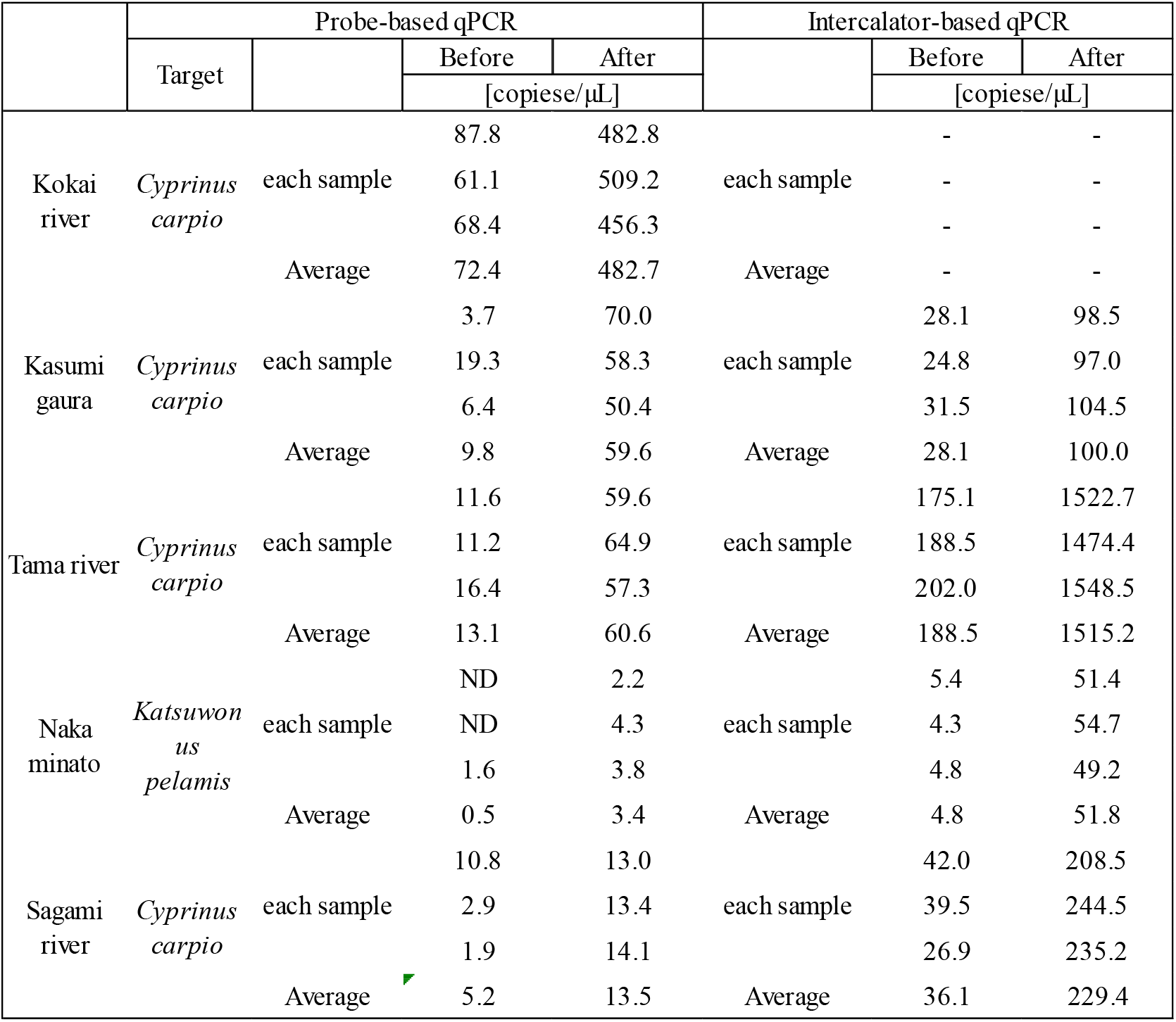
The results of probe-based qPCR and intercalator-based qPCR.

